# Accurate Detection of Proteins in Cryo-Electron Tomograms from Sparse Labels

**DOI:** 10.1101/2022.09.19.508602

**Authors:** Qinwen Huang, Ye Zhou, Hsuan-Fu Liu, Alberto Bartesaghi

**Affiliations:** Duke University, Durham NC 27708, USA

**Keywords:** Cryo-electron microscopy, cryo-electron tomography, 3D detection, positive-unlabeled training, contrastive learning

## Abstract

Cryo-electron tomography (CET) combined with sub-volume averaging (SVA), is currently the only imaging technique capable of determining the structure of proteins imaged inside cells at molecular resolution. To obtain high-resolution reconstructions, sub-volumes containing randomly distributed copies of the protein of interest need be identified, extracted and subjected to SVA, making accurate particle detection a critical step in the CET processing pipeline. Classical template-based methods have high false-positive rates due to the very low signal-to-noise ratios (SNR) typical of CET volumes, while more recent neural-network based detection algorithms require extensive labeling, are very slow to train and can take days to run. To address these issues, we propose a novel particle detection framework that uses positive-unlabeled learning and exploits the unique properties of 3D tomograms to improve detection performance. Our end-to-end framework is able to identify particles within minutes when trained using a single partially labeled tomogram. We conducted extensive validation experiments on two challenging CET datasets representing different experimental conditions, and observed more than 10% improvement in mAP and F1 scores compared to existing particle picking methods used in CET. Ultimately, the proposed framework will facilitate the structural analysis of challenging biomedical targets imaged within the native environment of cells.

## 1 Introduction

Cryo-electron tomography (CET) combined with sub-volume averaging (SVA) is currently the only imaging technique capable of determining the structure of proteins imaged inside cells at molecular resolution [8]. Bypassing the need for protein purification, CET allows determination of protein structures within their native context while also providing information on their distribution and partner interactions. Unlike cryo-EM single particle analysis (SPA) that requires sample purification and collects 2D projections of particles [2], CET can recover 3D information from proteins by recording a series of 2D images as the biological sample is rotated around a tilt axis (Figure 1). The sequence of 2D images, termed a *tiltseries*, is then aligned and used to calculate a 3D tomographic reconstruction or *tomogram* of the sample. A typical CET dataset usually contains between tens to a few hundred tomograms and each tomogram contains a few hundred copies of the same protein of interest. To obtain a single high-resolution structure, tens of thousands of sub-volumes containing randomly oriented and distributed copies of the protein of interest first need to be detected within tomograms in a process commonly referred to as *particle picking*. Sub-volumes are then extracted, aligned and combined in 3D using SVA, making the detection task critical for the downstream data processing. The low signal-to-noise ratios (SNR) characteristic of CET images, caused in part by the limited electron doses used during acquisition to prevent radiation damage of the biological samples, makes particle localization very challenging. High false-positive rates can prevent successful 3D reconstruction altogether due to the presence of confounding sub-volumes corresponding to noise, while high true-positive rates are desirable as they can lead to better denoising performance and increased resolution of the final reconstruction. For 2D SPA data, fully-supervised and semi-supervised learning-based particle picking methods can achieve good results due to the higher SNRs and availability of large annotated datasets [14]. In contrast, training of deep learning-based 3D CET particle picking algorithms remains impractical due to: (1) the lack of enough annotated CET datasets caused by the time consuming nature of doing manual labeling of 3D tomograms, and (2) the challenges imposed by the intrinsically lower SNR of tomographic projections and the effects caused by molecular crowding in native cellular environments. Recent efforts to tackle particle picking from CET tomograms using deep-learning only work on simulated datasets with full annotation, and use network architectures with millions of parameters that take days to train and hours to perform detection on a single tomogram, making their use impractical in real applications [11]. Currently, particle picking from CET tomograms remains a major bottleneck that has slowed down the reconstruction of high-resolution protein structures using SVA. To overcome the challenges in training deep learning-based particle identification models for CET, we propose a semi-supervised learning-based framework that only requires a few annotations on a single tomogram. The proposed method can be trained within minutes, which makes it suitable for a data-specific model. Specifically, our approach uses a positive unlabeled learning-based center localization module, allowing us to leverage information from both annotated and unlabeled data, effectively removing the burden of doing full data annotation. To enable better feature representation learning, we adopt a voxel-level contrastive learning module. The proposed module exploits both supervised and self-supervised contrastive learning and improves learned features. To validate our approach, we carried out extensive experiments on two challenging CET datasets acquired under different SNR conditions. We show that our method is able to outperform existing methods, while requiring smaller amounts of training data (less than 0.5% of the total data) and being time-efficient (under 10 minutes).

**Fig. 1:**
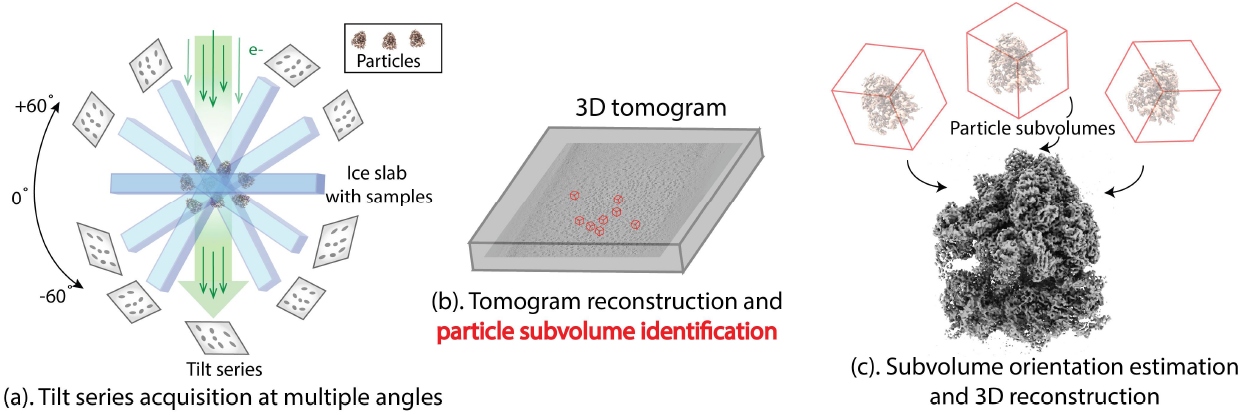
Overall processing pipeline for CET. (a) Projections of the protein sample are acquired at different angles by rotating the microscope stage using small tilt increments. (b) The acquired tilt-series are aligned and used to reconstruct 3D tomograms containing a few hundred sub-volumes which are identified and extracted. (c) Orientations of each extracted sub-volume are estimated followed by averaging to obtain the final high-resolution 3D reconstruction.

To summarize, the main contributions of this paper are:

1. We propose a 3D particle detection framework that achieves high localization accuracy with only a few annotations. The framework consists of two modules: (1) a positive unlabeled learning-based particle center localization module, and (2) a debiased voxel-level contrastive learning module. Both modules leverage information from annotated and unannotated data.
2. To the best of our knowledge, our work is the first to enable protein identification of hundreds of tomograms within minutes (training time included).
3. Through extensive experiments, we demonstrate that our framework is robust and performs well, even under the challenging SNR conditions of CET.

## 2 Related Work

### Object Detection and Applications in Cryo-Electron Microscopy (EM)

Neural network based object detection algorithms have been applied and shown promising results in various fields ranging from photography to medical imaging. Existing object detection approaches can be broadly divided into two categories: anchor-based and anchor-free. Representative examples of anchor-based methods include faster R-CNN [26], a region-based two-stage object detector, and YOLO [25] and SSD [21], which are one-stage object detectors. To address problems with class imbalance, the use of a focal loss term was proposed in [20]. Building on top of anchor-based methods, anchor-free methods were later introduced. This category includes FCOS [31], which performs per-pixel bounding box prediction, and CenterNet [36], which predicts bounding box location by estimating its center coordinates. Following the success of object detection algorithms based on neural networks, there have been multiple CNN-based particle picking algorithms for 2D cryo-EM SPA [13,1,23], including Topaz [3], a positive-unlabeled learning-based algorithm that learns to identify particles by minimizing both the supervised classification loss and the divergence between estimated empirical distribution and prior distribution; and crYOLO [32], a YOLO-based fully supervised particle detector. In contrast, there is very limited work on particle identification from 3D CET tomograms, where the most commonly used particle detection method is template matching [28]. Examples include recently proposed 3D-CNN based methods [11,22,34], and concurrent work, DeepPict [30], that uses a 2D-CNN for segmentation and a 3D-CNN for particle localization. These methods all require a large amount of annotated data and long training time.

### Positive Unlabeled Learning

PU learning can be broadly generalized into two categories: (1) two-step techniques that first identify reliable negative examples and learn based on labeled positives and reliable negatives; (2) class prior incorporation. The two-step techniques are similar to the teacher-student model that have been widely adopted in semi-supervised learning [33,27,16]. Class prior information can be incorporated in two ways: (1) the expected distribution of the classified unlabeled data should match the known prior distribution (this is a form of posterior regularization called GE criteria [9] and this approach is adopted by [3]), and (2) unbiased PU learning, where the unlabeled data is used as negatives while being properly down-weighted [18,24]. Based on unbiased PU learning, debiased instance level contrastive learning that takes sampling bias into account was also proposed [7].

### Contrastive Representation Learning

The goal of contrastive learning is to learn an embedding space in which similar sample pairs are close to each other while dissimilar pairs are far apart. Our proposed contrastive learning utilizes the InfoNCE loss, which has been adopted by many self-supervised contrastive learning frameworks such as simCLR [5] and MoCo [12]. In most self-supervised contrastive learning frameworks, heavy data augmentation, large batch size and hard negative sampling are crucial components. More recently, supervised contrastive learning [17] using InfoNCE was proposed and shown to be a generalization of triplet loss and N-pair loss. In addition to image-wise contrastive learning, there are pixel-wise contrastive learning frameworks used for image segmentation [35,4]. In our case, instead of using contrastive learning as a pre-training framework, we use it as a regularization component to the detection task.

## 3 Methodology

We first introduce the problem formulation of semi-supervised protein localization in crowded CET volumes and some special characteristics of CET data. We then give an overview of our proposed framework, which consists of two modules: positive unlabeled learning-based protein center localization, and voxel-level debiased contrastive feature learning. A detailed description of each module is then given. Finally, we present the overall training objective.

### 3.1 Characteristics of CET volumes

There are two important properties of CET tomograms, apart from their low SNR nature (Figure 3). First, due to the use of limited tilt angle ranges, the reconstructed tomograms contain a “missing-wedge” of information that distorts particle images due to lack of full orientation information. This distortion is especially obvious from the *Y* − *Z* view of the data which is perpendicular to the specimen plane. Second, there is data recurrence within each tomogram since a single tomogram usually contains up to a few hundred sub-volumes of the protein-of-interest. These copies are all present in different relative orientations (with respect to the missing wedge) and are distorted in different ways. Our particle detection technique leverages these unique properties of CET datasets.

### 3.2 Problem Formulation

The aim of semi-supervised protein localization in CET volumes is to obtain a model that is able to detect locations of proteins-of-interest in 3D tomograms, by learning from just a few annotated examples. A typical CET dataset 𝒟contains *j* tomograms {*T*_*i*_ ∈ *R*^*W* ×*H*×*D*^, *i* = 1, …, *j*}, with *j* ranging from tens to a few hundreds. A single tomogram *T*_*i*_ is scattered with a few hundreds to thousands of proteins. In the semi-supervised protein identification setting, in order to reduce the labor of manual labeling, the training set 𝒟_*tr*_ includes one tomogram *T*_*tr*_ with a few proteins annotated. The remaining tomograms are used as testing set 𝒟_*te*_. Unlike standard object detection algorithms which aim to produce bounding box locations of objects-of-interest, in most cryo-EM applications, center coordinates of proteins are the desired outputs for the particle detection task. Therefore, instead of outputting bounding box locations and sizes, inspired by [36], we aim to train the particle detector using 𝒟 ;_*tr*_ to produce a center point heatmap 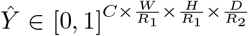 where *R*_*i*_ is the output stride and *C* is the number of protein species. The output stride down-samples the prediction by a factor *R*_*i*_ on each dimension. We set *C* = 1 as we only consider monodisperse samples and omit the dimension in the following sections. We extend the method used by [19] to generate the ground truth heatmap 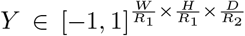 using the partially annotated tomogram. For each annotated center coordinate position *p* = (*x, y, z*), its downsampled equivalent is computed as 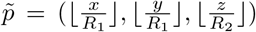. For each 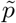 on *Y*, we apply a Gaussian kernel 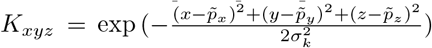 where *σ*_*k*_ is determined by the particle size [19]. The remaining unlabeled coordinates on *Y* have a value of −1.

### 3.3 Proposed Approach

#### Overview

As shown in Figure 2, our framework is composed of: (a) an encoderdecoder feature extraction backbone, (b) a protein center localization module, and (c) a voxel-level contrastive feature learning module. We used a fully convolutional architecture for the backbone and since the input training tomogram is only partially labeled, we incorporated a positive unlabeled learning-based strategy for both the localization and contrastive learning modules.

**Fig. 2:**
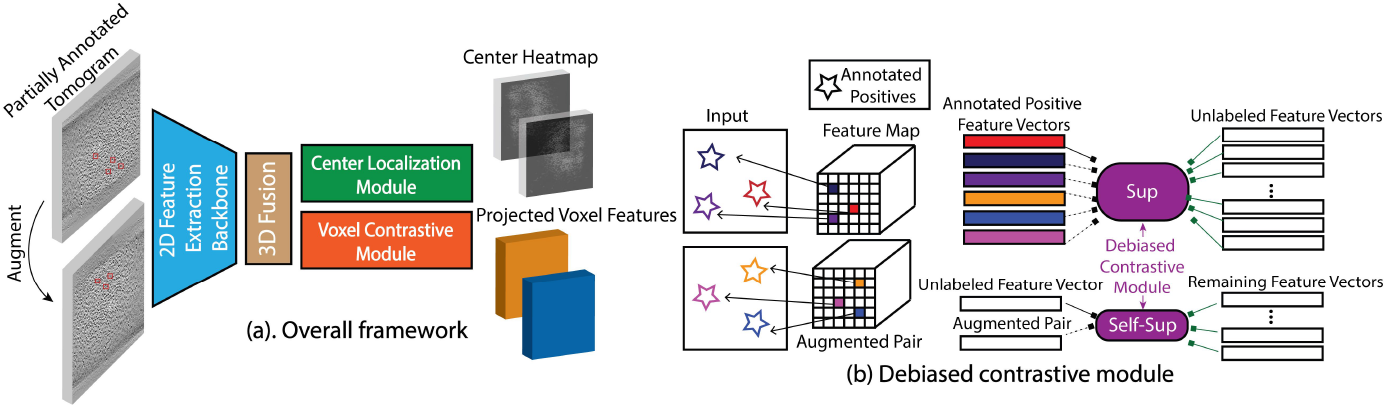
(a) Proposed framework for 3D protein detection. We use a combination of 2D and 3D convolutional layers. The 2D CNN-based feature extractor follows a encoder-decoder architecture. The feature extractor is applied to each slice of the tomogram. The extracted features are then fused together through 3D convolutional layers. The fused 3D features are used for: (1) center coordinate heatmap prediction, and (2) voxel-level contrastive learning. During inference, only the heatmap is used for particle identification. **(b) Voxel-level debiased contrastive learning module**. For illustration purposes, we use a 2D image with stars in different orientations. These stars represent naturally augmented positive pairs. Feature vectors at the star locations encode information about the location of the objects and the feature vectors serve as input to our contrastive learning module.

#### Feature extraction backbone

Even though the input is a 3D tomogram, our network is composed of mostly 2D convolutional layers. 3D convolutional layers are only applied in the last two layers. Essentially, the network first extracts features of each slice independently and then merges the extracted 2D features into 3D at the final layers. The combination of 2D and 3D layers is inspired by the actual manual particle picking process: to identify a particle, the *X*−*Y* view of each slice is inspected most carefully (as it contains no distortions), while the *X*− *Z* and *Y*− *Z* views only provide very limited information (due to the heavy missing-wedge distortions, Figure 3). This architecture design has two advantages: first, since 3D information is only considered during the final layers, it can reduce the missing-wedge effect; second, the resulting architecture has fewer parameters than a pure 3D CNN, greatly reducing memory requirements and running time. We provide more details in the supplementary material.

**Fig. 3.**
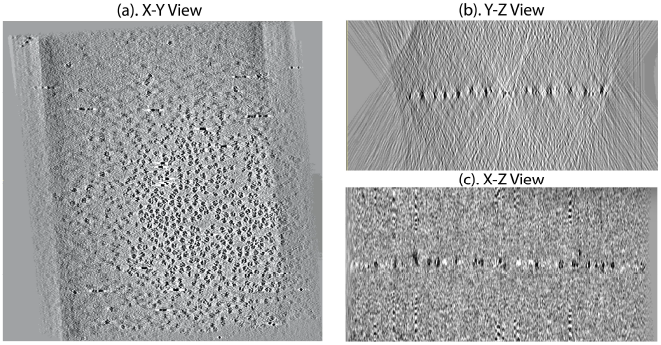
An example CET slice from all three views. (a) X-Y view. (b) Y-Z view. (c) X-Z view. X-Y view provides most amount of useful information for particle identification.

#### Protein center localization module

For the input tomogram *T* and its output heatmap *Ŷ*, protein localization can be viewed as a per-voxel classification problem such that each voxel *v*_*i,j,k*_ at position (*i, j, k*) is the input and the corresponding *ŷ*_*i,j,k*_ ∈ [0, 1] is the classification output.

##### Positive Negative (PN) Learning

Denote *p*(*v*) as the underlying data distribution from which *v*_*i,j,k*_ is sampled, *p*(*v*) can be decomposed as follows:

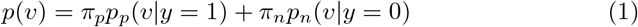

where *p*_*p*_(*v*|*y* = 1) is the positive class conditional probability of protein voxels, *p*_*n*_(*v*|*y* = 0) is the negative class conditional probability of background voxels, and *π*_*p*_ and *π*_*n*_ are the class prior probabilities. Underscripts *n, p, u* denote negative, positive and unlabeled, respectively. Denote *g* : ℝ ^*d*^→ ℝ, an arbitrary classifier that can be parameterized by a neural network, *l*(*g*(*v*) = *ŷ, y*) being the loss between model outputs *ŷ* and ground truth *y*. When all the voxels are labeled, this is essentially a binary classification problem that can be optimized using a standard PN learning approach with the following risk minimization:

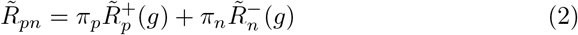

where 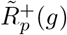 is the mean positive loss 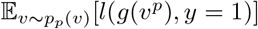 and can be estimated as 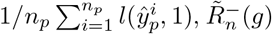 is the mean negative loss 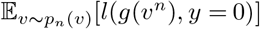 and can be estimated as 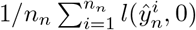 *n*_*p*_ and *n*_*n*_ are the number of positive and negative voxels.

##### |Positive Unlabeled (PU) Learning

When only a few positive voxels are labeled and the remainder of the data is unlabeled, we re-formulate the problem into the PU setting: the positive labeled voxels are sampled from *p*_*p*_(*v* | *y* = 1) and the remaining unlabeled voxels are sampled from *p*(*v*). As shown in [24], by rearranging Equation 1 and 2, we obtain *π*_*n*_*p*_*n*_(*v*) = *p*(*v*) − *π*_*p*_*p*_*p*_(*v*) and 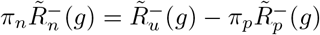. We therefore rewrite the risk minimization as:

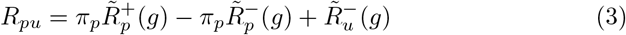

with 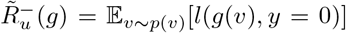 and 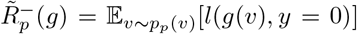. In order to prevent overfitting in Equation 3, we adopted the non-negative risk estimation as in [18]:

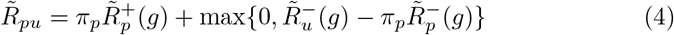

##### Soft Positives and True Positives

Since the ground-truth heatmap is splatted with Gaussian kernels, the labels are not strictly binary. Positive labels are split into two groups: true positives (tp) where *y*_*i,j,k*_ = 1, which is the center of each Gaussian kernel (protein center), and soft positives (sp) where 0 < *y*_*i,j,k*_ < 1 (voxels that are close to the center). Unlabeled voxels are labeled as 1. With this, the positive distribution *p*_*p*_(*v*) and positive associated losses 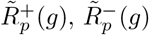 are decomposed into:

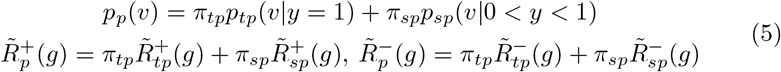

We adopt voxel-wise logistic regression with focal loss for *l*(*g*(*v*), *y*). Specifically, we have:

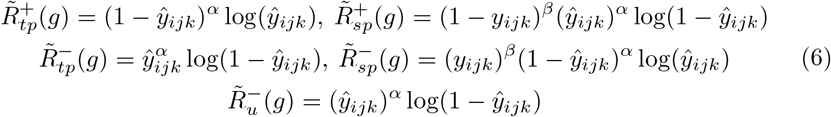

where *α, β* are the focal loss parameters and we use *α* = 2, *β* = 4. By combining Equation 4, 5 and 6, we obtain the final minimization:

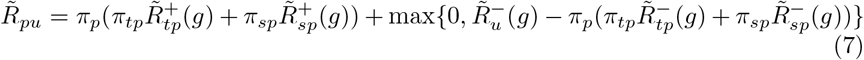

#### Voxel level debiased contrastive learning module

For the input tomogram *T* ∈ R^*W* ×*H*×*D*^ and its augmented pair 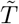, denote the output from the feature extraction backbone as 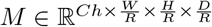 and the augmented pair 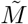. *M* and 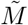 are used to generate: (1) the output heatmap *Ŷ*and its augmented pair *Ŷ*, and (2) the projected feature map *F* and 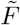 As suggested in [5], instead of using *M*, it is beneficial to map the representations to a new space through a projection head composed of 1× 1× 1 convolutional layer where a contrastive loss is applied. Denote *m*_*i,j,k*_ ∈ ℝ^*Ch*^ and 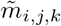 as the feature vector at the (*i, j, k*) position of the feature map *M* and its augmented counterpart. There exists a total of 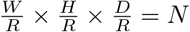 such vectors. Each of these feature vectors is responsible for predicting *ŷ*_*i,j,k*_. If *y*_*i,j,k*_ = 1, *m*_*i,j,k*_ and its projection *f*_*i,j,k*_ should encode particle-related features. For a partially annotated *T*, the voxel-level feature vectors *f* can be separated into positive and unlabeled classes. Therefore, the voxel-level contrastive loss is composed of: (1) positive supervised debiased contrastive, and (2) unlabeled self-supervised debiased contrastive terms.

##### Positive supervised debiased contrastive loss

Denote 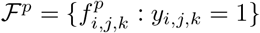 as the set of positive feature vectors obtained from *n*_*p*_ annotated proteins and its augmented counterpart 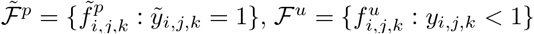 as the set of unlabeled (including the soft positives) feature vectors with a total of *n*_*n*_. Since each tomogram contains up to a few hundred sub-volumes of the protein of interest, and each of these sub-volumes are the same protein with different relative orientations and are distorted in different ways, for a feature vector 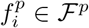 the remaining 2*n*_*p*_ − 1 feature vectors 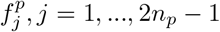 in ℱ^*p*^ and 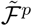 can be treated as its naturally augmented pair. Unlabeled feature vectors 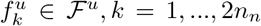which includes the augmented unlabeled features are treated as negatives. However, since the unlabeled setℱ ^*u*^ can contain positive feature vectors, the naive supervised contrastive loss as proposed in [17] will be biased. We therefore adopt a modified debiased supervised contrastive loss based on [7]:

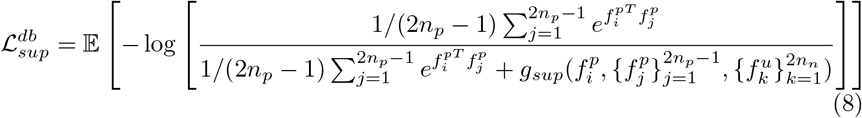

where the second term in the denominator is:

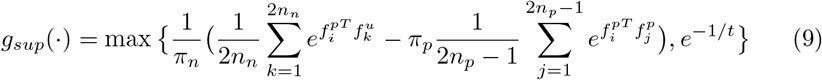

with *π*_*n*_ and *π*_*p*_ being the class prior probabilities and *t* the temperature.

##### Unlabeled self-supervised debiased contrastive loss

For the unlabeled feature vector 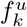, the only known positive is its augmented pair 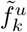 and the remaining vectors are treated as negatives. Denote 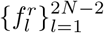 as the set of remaining vectors. The resulting contrastive loss for an unlabeled feature vector is:

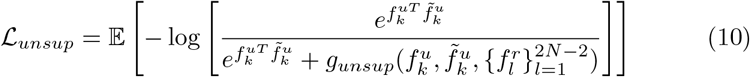

It should be noted that *g*(·) involves class prior probabilities, however, the actual class of the unlabeled feature vectors is unknown. Therefore, we used the output probabilities from *Ŷ* and

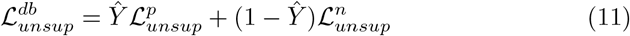

where for 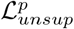, the denominator 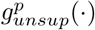 is:

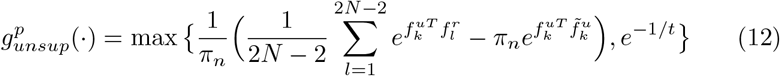

and for 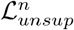, the denominator 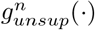 is:

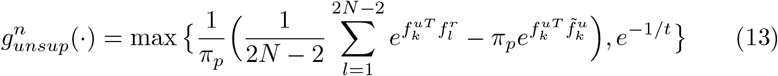

#### Overall Training Objective

In addition, we added a consistency regularization loss for the output heatmap *Ŷ* and its augmented version *Ŷ* such that the probability of a voxel containing a protein should be invariant to augmentations:

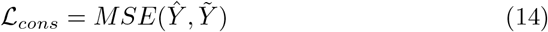

The final training objective is:

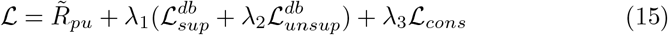

where *λ*_1_ is the weight of the total contrastive module, *λ*_2_ is the weight of the unsupervised contrastive loss, and *λ*_3_ is the weight of consistency regularization. The resulting loss serves two purposes: 1) the contrastive term maximizes similarities for encoded features belonging to the same group (particle and background) and minimizes such similarities if features are from different groups, and 2) the heatmap loss term forces predicted particle probabilities to be higher when they are closer to the true center location. To remove duplicate predictions, non-max suppression is applied to the predicted heatmap using 3D max-pooling.

## 4 Validation Experiments

We evaluate the performance of our algorithm on two real CET datasets. For each dataset, we evaluated the performance when 5%, 10%, 30%, 50% and 70% of the data from a single tomogram is annotated. We perform ablation studies over the contrastive and positive unlabeled learning components. Performance is measured using mean average precision (mAP) scores calculated against manually labeled particle locations.

### 4.1 Datasets

We evaluate our method on two publicly available CET datasets from the electron microscopy public image archive (EMPIAR) [14]: EMPIAR-10304 [10] and EMPIAR-10499 [29]. These datasets represent the two most common types of CET biological samples, more details are included in supplementary material.

### 4.2 Experimental Setup

#### Implementation Details

We initialize the network using weights obtained from a trained network that identifies whether a slice contains particles. However, random initialization of the network is able to achieve similar performance. During training, instead of using the whole tomogram, we cropped sub-tomograms of size 64 ×64 × 5 as input to the network in batches of 2. Training time is thus independent of input size. Inference is performed on the entire tomogram. The network is implemented with PyTorch and trained/tested on an NVIDIA Tesla V100 GPU. The proposed framework is trained in an end-to-end manner using Adam optimizer with default parameter values and an initial learning rate of 0.001. We decrease the learning rate by a factor of 10 every 200 iterations. Training takes around 3 to 5 minutes for 600 iterations and inference on each full tomogram takes less than a second. In all our experiments, we trained the network for 600 iterations. We used experimentally determined values: *λ*_1_ = 0.1, *λ*_2_ = 0.5 and *λ*_1_ = 0.1. For EMPIAR-10304, we used *π*_*p*_ = 0.6, and *t* = 0.07. For EMPIAR-10499, we used *π*_*p*_ = 0.1, and *t* = 0.02.

#### Evaluation Metrics

We use mean average precision (mAP) scores calculated against manually labeled particle locations for evaluation. To account for small variations in the detected particle centers, instead of looking at a single pixel, we also look at pixels located within a certain radius from the center. If the detected particle position is within a certain radius of a ground truth particle position, it is considered as a true positive match. Similarly, if there is no ground truth particle within a certain radius of a detected particle position, it is considered as a false positive. We use radius values of 2 and 5 and denote the corresponding mAP values as *mAP*_*r*2_ and *mAP*_*r*5_.

#### Baseline Methods

We compare our method with one conventional CET particle detection method, template matching, and one recently developed deep learning-based method, crYOLO-3D [32]. Even though there are many available deep-learning based particle picking methods for 2D SPA cryo-EM, there are only a few methods for 3D CET. crYOLO-3D is the only deep learning-based algorithm that is available to use. Template matching is implemented using the EMAN2 package [28]. We use a low-pass filtered ribosome reconstruction as template. Even though crYOLO-3D is termed as “3D” picking, it is really a 2D picking method. For each 3D tomogram input, crYOLO-3D performs per-slice particle detection using a pretrained 2D model. The detected 2D coordinates on each slice are combined into 3D coordinates through a tracking algorithm. The pretrained model is trained using 43 fully labeled datasets with more than 44,000 labeled particles. We use their official software available online^1^ and fine tune the pretrained weights with images and labels from our training samples (Note: we had to convert 3D labels to 2D (slice-level) for crYOLO training, and therefore we used 360 annotations for EMPIAR-10304 and 1100 annotations for EMPIAR-10499). For the baseline methods, we are only able to obtain precision and recall values, as their implementation does not output detection scores and we can only select a cut-off threshold. Therefore, for comparing with our framework, we also calculated the corresponding precision-recall score using the same threshold (0.25).

### 4.3 Results

In Table 2, we show mAP_*r*2_ and mAP_*r*5_ scores for both datasets obtained using our approach. We show qualitative visualization of detection results on selected slices of tomograms from each dataset in Figure 4. To improve particle visibility, we use averages of multiple slices instead of a single slice. Our method is able to outperform the two baseline methods by a significant margin, as shown in Table 1. We also show that when trained only using a very small amount of annotated data (5% and 10%), our approach is still able to obtain satisfactory results. When more annotations are available for training, performance improves (especially from 5% to 30%). Template matching from EMAN2 tends to pick up more false positives and is less accurate in identifying the true center of the particle. It tends to miss more particles when SNR is lower. While crYOLO3D is more robust to noise compared to template matching, it still results in many missed particles. In addition, it performs poorly on the extremely crowded dataset (EMPIAR-10304) even though the SNR is much higher. This is because crYOLO-3D is actually a 2D particle detector. Instead of processing the entire tomogram as one 3D volume, the inputs are individual 2D slices: it performs particle detection on each slice first. The 2D coordinate outputs on each slice are then merged in the post-processing step (multi-slice tracing) into 3D coordinates. During the merging step, the actual 3D tomogram is not used, only the 2D coordinates are used as inputs to the post-processing step. When particles are crowded, tracing fails and the subsequent detection also fails. Since our algorithm uses the whole 3D tomogram as input, we are able to avoid these problems. Our method is also able to avoid contamination areas, as features corresponding to these areas are learned to be distinct from features characteristic of true particles.

**Fig. 4:**
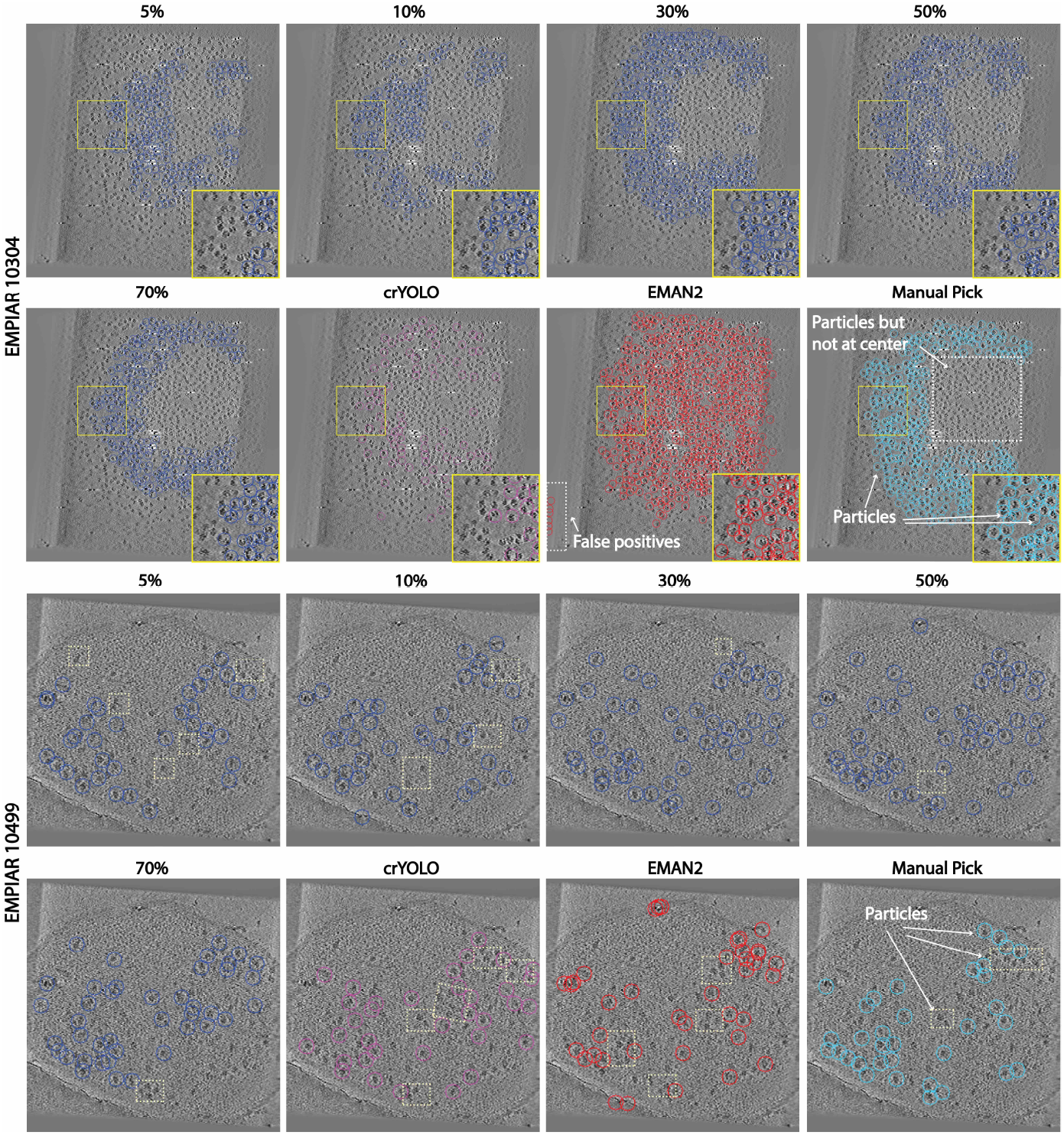
Detection results on selected tomograhic slices from EMPIAR-10304 and EMPIAR-10499. Top two rows: Slice averages from tomograms of EMPIAR-10304. Zoomed-in views are provided at the bottom right. Bottom two rows: Slices averages from tomograms of EMPIAR-10499. We show detected particles trained using 5%, 10%, 30%, 50% and 70% of particles annotated on a single tomogram. We also show particle detection results using crYOLO-3D and EMAN2. Our method is able to detect more particles with highere precision. As more data annotation is available, detection performance increases (especially from 5% to 50%). EMAN2 tends to pick up more false positives. Note that manual picked results are not necessarily ground truth, as it is possible to miss several particles during manual picking (as shown). We highlighted several missed particles regions in light yellow.

**Table 1:** Precision, Recall and F1 scores for our proposed method and baseline methods.

| Method | EMPIAR 10304 |  |  | EMPIAR 10499 |  |  |
| --- | --- | --- | --- | --- | --- | --- |
|  | Precision | Recall | F1 | Precision | Recall | F1 |
| 5% | 80.4 | 59.2 | 68.2 | 49.6 | 58.1 | 53.5 |
| 10% | 81.8 | 53.6 | 64.7 | 50.1 | 58.2 | 53.8 |
| 30% | 79.5 | 76.5 | 78.0 | 55.9 | 60.3 | 58.0 |
| 50% | 83.8 | 72.1 | 77.5 | 53.0 | 65.1 | 58.4 |
| 70% | 82.3 | 77.6 | 80.0 | 54.9 | 66.7 | 60.2 |
| crYOLO | 47.6 | 14.6 | 22.4 | 47.8 | 56.8 | 52.0 |
| EMAN2 | 58.9 | 65.5 | 62.0 | 26.1 | 55.3 | 35.5 |

<sup>1</sup> https://cryolo.readthedocs.io/en/stable/

**Table 2:** Particle detection results obtained using different levels of annotations available for training. *First row* : mAP scores for our proposed method with both positive unlabeled center localization module and debiased voxel-level contrastive learning module. *Second row* : mAP scores for our proposed method without the contrastive learning module. *Third row* : Our proposed method without positive unlabeled learning and debiased contrastive learning

| EMPIAR-10304 |  |  |  |  |  |  |  |  |  |  |
| --- | --- | --- | --- | --- | --- | --- | --- | --- | --- | --- |
| mAP | 5% |  | 10% |  | 30% |  | 50% |  | 70% |  |
|  | r5 | r2 | r5 | r2 | r5 | r2 | r5 | r2 | r5 | r2 |
| Ours | **54.5** | **44.0** | **57.0** | **45.2** | **67.9** | **56.8** | **71.5** | **62.0** | **72.1** | **62.5** |
| Ours no CR | 53.0 | 42.1 | 54.1 | 43.2 | 65.1 | 55.7 | 67.5 | 58.1 | 65.8 | 57.4 |
| Ours no PU | 32.2 | 14.1 | 38.3 | 22.9 | 41.5 | 15.4 | 40.2 | 20.0 | 41.1 | 23.1 |

**Table 2: Particle detection results obtained using different levels of annotations available for training.**
| EMPIAR-10499 |  |  |  |  |  |  |  |  |  |  |
| --- | --- | --- | --- | --- | --- | --- | --- | --- | --- | --- |
| mAP | 5% |  | 10% |  | 30% |  | 50% |  | 70% |  |
|  | r5 | r2 | r5 | r2 | r5 | r2 | r5 | r2 | r5 | r2 |
| Ours | **31.1** | **22.0** | **33.2** | **22.4** | **40.1** | **29.8** | **41.3** | **30.2** | **42.5** | **31.3** |
| Ours no CR | 24.2 | 14.1 | 24.3 | 14.2 | 33.9 | 24.0 | 31.8 | 19.1 | 34.5 | 24.1 |
| Ours no PU | 11.4 | 2.1 | 9.4 | 1.7 | 16.2 | 4.1 | 14.8 | 3.5 | 20.0 | 9.2 |

In terms of running time and level of annotation, template matching takes 2-5 minutes and a template is required. When a template is not available, a Gaussian blob is used, which results in even higher false positive rates. crYOLO-3D takes a total of 30-40 minutes to run including model fine-tuning. Our proposed method takes 5-10 minutes to run and requires a minimum of 20-50 labeled particles.

#### Ablation Studies

To evaluate the effectiveness of our proposed method we perform ablations studies on: (1) voxel-level contrastive learning module, and (2) positive unlabeled learning in two modules. For (1), we remove the proposed voxel-level contrastive module and the corresponding term in the loss function. As shown in Table 2, without this module, the performance degrades, especially under lower SNR scenarios. This shows that our proposed module improves the feature learning of input tomograms and in turn facilitates the detection of particles when only limited amount of training samples are provided. In (2), for the center localization module, we treat all unlabeled regions as negatives and adopted a standard focal loss as in [36]; for the contrastive module, we adopted a combination of supervised contrastive loss as in [17] for labeled regions and self-supervised InfoNCE for unlabeled regions. As shown in Table 2, the detection outcome decreases significantly without positive unlabeled learning, which implies the importance of debiasing when there is lack of annotated data. We also evaluated the effect of feature extraction backbone choices on detection outcomes. For this, we looked at: (1) 2D convolution vs. 3D convolution, and (2) depth of the network. For (1), even though for volumetric data 3D convolution-based architectures are more commonly used [6,15], due to the unique properties of CET data and the lack of training data, we experimentally found out that full 3D convolution-based architectures (3D ResNet and UNet) failed to learn any useful information, which is why we did not include their corresponding results in this section. For (2), as objects-of-interest in CET are small, increasing the depth of networks (which increases the receptive field size) can actually worsen performance. We include more details in the supplementary material.

#### Limitations

The main limitation of our proposed method is the necessary knowledge of the class prior probabilities *π*. For crowded samples like EMPIAR-10304, if we use a very small positive prior such as 0.1, the trained model tends to produce more false negatives. On the contrary, for less crowded samples like EMPIAR-10499, if we use a large positive prior, more false positives get identified. Therefore, a reliable estimation of *π* is required. Such estimation can be obtained by visually inspecting the tomogram when doing annotations. In addition, the performance of our method is limited under very low SNR levels.

## 5 Conclusion

We propose a novel 3D particle detection framework that enables accurate localization of proteins from CET datasets within minutes when trained using a small amount of labeled data. By leveraging the internal data statistics of CET tomograms, we design a novel architecture for 3D particle identification that incorporates positive unlabeled and contrastive learning. Extensive experiments demonstrate that our proposed framework achieves superior performance on real cryo-ET datasets compared to previous methods. The proposed framework will expedite the current cryo-ET data processing pipeline and facilitate the structural analysis of challenging biomedical targets imaged within cells.

## Acknowledgements

This study utilized the computational resources offered by Duke Research Computing (http://rc.duke.edu). We thank C. Kneifel, K. Kilroy, M. Newton, V. Orlikowski, T. Milledge and D. Lane from the Duke Office of Information Technology and Research Computing for providing assistance with the computing environment. This work was supported by a Visual Proteomics Imaging grant from the Chan Zuckerberg Initiative (CZI) to AB.

## Footnotes

1 https://cryolo.readthedocs.io/en/stable/

