## Supplementary Information for "Accurate Detection of Proteins in Cryo-Electron Tomograms from Sparse Labels"

### 1 Architecture

We tested our proposed framework using: 1) ResNet-based architecture, and 2) UNet-based architecture. Detailed architectures are shown in the following. The main difference between these two architectures is the skip connection. Through skip connection, the decoder layers have access to feature maps at higher levels, thereby preserving finer detail information.

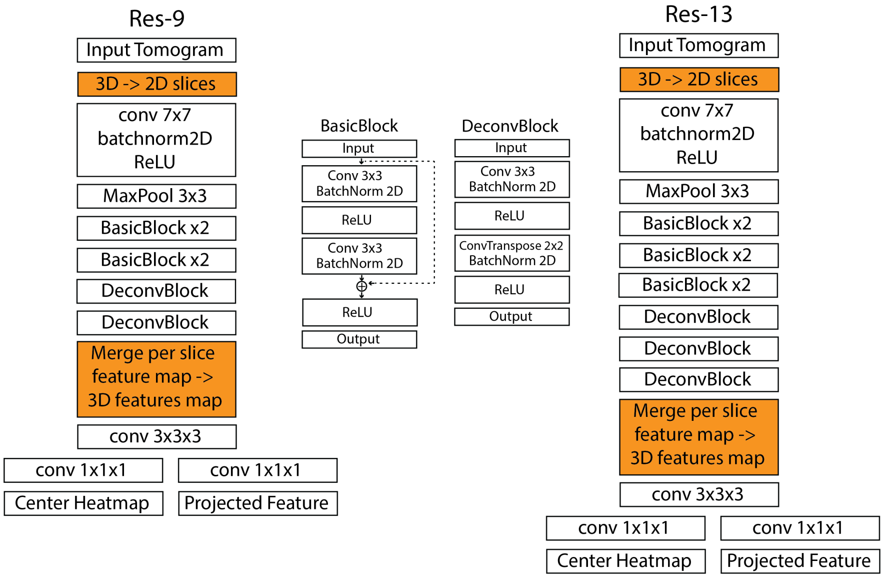


**Fig. 1: ResNet-based architecture.** Res-9 includes 4 basic blocks and Res-13 includes 5 basic blocks for encoder. As shown, most convolutional operations are performed in 2D, with the final layers being 3D. Res-9 encoder performs 8x downsampling and Res-13 encoder performs 16x downsampling.

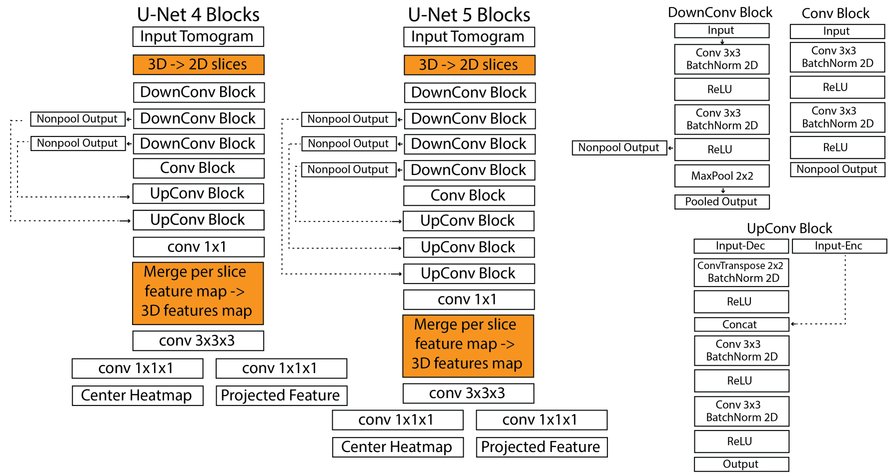


**Fig. 2: UNet-based architecture.** UNet-4 encoder includes 4 blocks and UNet-5 encoder includes 5 blocks. As shown, most convolutional operations are performed in 2D, with the final layers being 3D. UNet-4 encoder performs 8x downsampling and UNet-5 encoder performs 16x downsampling.

### 2 Experimental Dataset

*EMPIAR-10304* This is a purified ribosome dataset that includes 12 tilt-series taken between $-60$ to $+60$ degrees of tilting. 12 tomograms of size $512 \times 512 \times 256$ were reconstructed using standard CET protocols [1]. To reduce input size, we merge every two $X - Y$ slices into one slice and obtained tomograms of size $512 \times 512 \times 128$. The predicted heatmap $\hat{Y}$ is of size $256 \times 256 \times 128$. Since this dataset is from a purified protein sample, ice thickness is relatively thin resulting in relatively high SNR and high visibility of particles. Each tomogram contains around 800-1000 particles. We used one partially annotated tomogram for training. 5% of labeling corresponds to a total of 45 particles.

*EMPIAR-10499* This is a ribosome dataset from native M. pneumoniae cells treated with chloramphenicol. The dataset contains 65 tilt-series taken from $-60$ to $+60$ degrees and tomograms of size $512 \times 512 \times 256$ are reconstructed. Ribosomes in this dataset are imaged within their native environment, resulting in thick ice and very low SNR and particle visibility. To reduce input size and improve SNR, we downsampled the tomograms in each dimension by a factor of two, resulting in volumes of size $256 \times 256 \times 128$. The predicted heatmap $\hat{Y}$ has size $128 \times 128 \times 128$. Tomograms from this native dataset have lower particle concentrations, each containing between 100 to 300 particles. We use one partially labeled tomogram for training. 5% of the labeling corresponds to a total 12 particles.

### 3 Experimental results with different architectures

Detection results obtained using different architectures are shown in Table 1. Specifically, we investigated the effect of depth of the network on detection performance. As shown, increasing depth does not necessarily improve the performance. On the contrary, if no skip connections exist, the performance worsens as depth increases (Res-9 vs. Res-13). This can be explained by the fact that as depth increases, the size of the receptive field also increases, and for smaller sized objects like particles, an increase in receptive field does not add useful information to their corresponding feature vectors. Skip connections can alleviate the problem by concatenating feature maps obtained at higher levels (which includes finer detail information since receptive field is smaller for higher levels) to the decoding layers. Overall, shallower network depths give better detection performance.

**Table 1: Particle detection results obtained using different architectures.**

| | **EMPIAR-10304** | | | | | | | | | |
|---|---|---|---|---|---|---|---|---|---|---|
| | 5% | | 10% | | 30% | | 50% | | 70% | |
| mAP | .5 | .75 | .5 | .75 | .5 | .75 | .5 | .75 | .5 | .75 |
| Res-9 | 54.5 | 44.0 | 57.0 | 45.2 | 67.9 | 56.8 | 71.5 | 62.0 | 72.1 | 62.5 |
| Res-13 | 35.5 | 12.6 | 43.9 | 14.4 | 47.3 | 11.2 | 44.6 | 17.2 | 41.9 | 17.9 |
| UNet-4 | 65.3 | 55.4 | 72.5 | 64.0 | 76.2 | 71.5 | 76.6 | 71.9 | 76.8 | 72.0 |
| UNet-5 | 67.2 | 57.9 | 66.8 | 57.9 | 70.2 | 65.0 | 78.9 | 69.2 | 79.0 | 73.1 |
| | **EMPIAR-10499** | | | | | | | | | |
| | 5% | | 10% | | 30% | | 50% | | 70% | |
| mAP | .5 | .75 | .5 | .75 | .5 | .75 | .5 | .75 | .5 | .75 |
| Res-9 | 31.1 | 22.0 | 33.2 | 22.4 | 40.1 | 29.8 | 41.3 | 30.2 | 42.5 | 31.3 |
| Res-13 | 16.5 | 5.2 | 19.1 | 6.1 | 19.3 | 6.2 | 22.1 | 12.9 | 24.9 | 13.1 |
| UNet-4 | 31.4 | 22.9 | 30.5 | 19.6 | 31.7 | 22.6 | 31.6 | 23.8 | 32.3 | 24.1 |
| UNet-5 | 28.0 | 17.1 | 29.1 | 18.2 | 27.9 | 17.0 | 28.2 | 17.5 | 27.6 | 18.1 |

### 4 Experimental results with consistency regularization

We also investigated how consistency regularization affects the performance of the proposed particle detector. As shown in Table 2, consistency regularization is able to improve the performance when fewer data annotations are available. When more labeled data is available, the benefit of consistency regularization is less significant.

Table 2: Effect of consistency regularization on particle detection

| | EMPIAR-10304 | | | | | | | | | |
|---|---|---|---|---|---|---|---|---|---|---|
| | 5% | | 10% | | 30% | | 50% | | 70% | |
| mAP | .5 | .75 | .5 | .75 | .5 | .75 | .5 | .75 | .5 | .75 |
| Res-9 | 54.5 | 44.0 | 57.0 | 45.2 | 67.9 | 56.8 | 71.5 | 62.0 | 72.1 | 62.5 |
| Res-9 no consis | 48.9 | 39.9 | 49.4 | 40.8 | 64.8 | 52.6 | 70.6 | 61.3 | 71.0 | 62.8 |
| UNet-4 | 65.3 | 55.4 | 72.5 | 64.0 | 76.2 | 71.5 | 76.6 | 71.9 | 76.8 | 72.0 |
| UNet-4 no consis | 64.1 | 54.0 | 70.8 | 60.9 | 73.6 | 68.5 | 75.6 | 69.7 | 76.4 | 71.6 |
| | EMPIAR-10499 | | | | | | | | | |
| | 5% | | 10% | | 30% | | 50% | | 70% | |
| mAP | .5 | .75 | .5 | .75 | .5 | .75 | .5 | .75 | .5 | .75 |
| Res-9 | 31.1 | 22.0 | 33.2 | 22.4 | 40.1 | 29.8 | 41.3 | 30.2 | 42.5 | 31.3 |
| Res-9 no consis | 30.4 | 21.5 | 29.7 | 21.4 | 38.2 | 29.0 | 40.3 | 30.8 | 40.8 | 31.0 |
| UNet-4 | 31.4 | 22.9 | 30.5 | 19.6 | 31.7 | 22.6 | 31.6 | 23.8 | 32.3 | 24.1 |
| UNet-4 no consis | 30.7 | 21.8 | 30.8 | 21.9 | 31.5 | 22.0 | 31.9 | 21.3 | 33.7 | 25.6 |
